# A specific set of heterogeneous native interactions yields efficient knotting in protein folding

**DOI:** 10.1101/2021.05.19.444793

**Authors:** João NC Especial, Patrícia FN Faísca

**Affiliations:** Departamento de Física and Faculdade de Ciências, Universidade de Lisboa, Campo Grande, Ed. C8, Lisboa, Portugal; BioISI - Biosystems and Integrative Sciences Institute, Faculdade de Ciências, Universidade de Lisboa, Campo Grande, Ed. C8, Lisboa, Portugal

**Keywords:** knotted proteins, protein folding, lattice model, Monte Carlo

## Abstract

Native interactions are crucial for folding, and non-native interactions appear to be critical for efficiently knotting proteins. Therefore, it is important to understand both their roles in the folding of knotted proteins. It has been proposed that non-native interactions drive the correct order of contact formation, which is essential to avoid backtracking and efficiently self-tie. In this study we ask if non-native interactions are strictly necessary to tangle a protein, or if the correct order of contact formation can be assured by a specific set of native, but otherwise heterogeneous, interactions. In order to address this problem we conducted extensive Monte Carlo simulations of lattice models of proteinlike sequences designed to fold into a pre-selected knotted conformation embedding a trefoil knot. We were able to identify a specific set of heterogeneous native interactions that drives efficient knotting, and is able to fold the protein when combined with the remaining native interactions modeled as homogeneous. This specific set of heterogeneous native interactions is strictly enough to efficiently self-tie. A distinctive feature of these native interactions is that they do not backtrack, because their energies ensure the correct order of contact formation. Furthermore, they stabilize a knotted intermediate state, which is enroute to the native structure. Our results thus show that - at least in the context of the adopted model - non-native interactions are not necessary to knot a protein. However, when they are taken into account into protein energetics it is possible to find specific, non-local non-native interactions that operate as a scaffold that assists the knotting step.

## Introduction

Knotted proteins embed a physical (i.e. open) knot in their native structure. The development of knot detection methods and loop closure procedures allowed determination of the fraction of knotted proteins in the protein data bank (PDB), as well as their knot type^1^. Knotted proteins exhibit most frequently the trefoil (or 3_1_) knot, but there are also proteins with 4_1_, 5_2_, and even one protein with the so-called Stevedore’s knot that comprises 6 crossings^2^. Depending on its location within the polypeptide chain the knot can be classified as shallow (when removing just a few residues from one of the termini is enough to untangle the fold) or deep.

These topologically complex proteins have attracted considerable attention and there is a large and growing body of experimental and theoretical work dedicated to understand their folding and knotting mechanisms^3–13^, their folding/unfolding properties^14–18^, their biological role^19–21^, and how they evolved^22–24^, just to mention a few examples.

Tracing and characterizing the folding pathway(s) is crucial to understanding the folding process of a globular protein. However, most standardly used experimental methods cannot yet adequately resolve the folding pathway with atomic detail and hence one must resort to molecular simulation in order to gain insights towards this goal. So far, several computational models based on protein representations with different levels of coarse graining, ranging from simple lattice to full atomistic representations, combined with different force fields and sampling methods, have been used to explore different aspects of the folding process of tangled proteins^25,26^.

Shakhnovich and co-workers carried out the first computational study^3^ that provided the very first vistas on the knotting mechanism of protein YibK (PDB ID: 1j85), which embeds a deep trefoil knot located 77 residues away from the N-terminus and 39 residues away from the C-terminus. They combined Langevin Dynamics simulations with a C-alpha representation to find that knotting occurs primarily in compact, near native conformations (with 80% of its native contacts formed), where part of the polypeptide chain is arranged into the so-called knotting loop, which is threaded by the C-terminus; the threading step occurs either directly (i.e. without bending) as observed for the vast majority of trajectories, or through an intermediate conformation in which the terminus arranges itself into an hairpin, and first gives rise to a pseudo-knotted conformation called a slipknot^58^. Interestingly, the authors reported a non-negligible probability of observing the formation of the knot in less structurally consolidated conformations (with only 20% of their native contacts formed). The most striking conclusion of this study concerns the energetics of the knotting step. Indeed, it was reported that when only native interactions (i.e. interactions that are present in the native structure) are considered in protein energetics (as in the native centric Gō models^27^), the conformations are too compact to allow for threading events, and none of the 100 attempted folding runs was successful. The authors reported that non-native interactions were essential to observe 100% folding efficiency (i.e. all attempted folding trajectories leading to native conformations). In particular, it was found that a specific set of ad-hoc non-native attractive interactions established between a stretch of residues within the knotting loop (86-93), and residues pertaining to the C-terminus assist the threading step.

A subsequent study by Faccioli and co-workers^28^ investigated the folding and knotting processes of AOTCase (PDB ID: 2g68), a 332 residue long protein embedding a deep trefoil knot in its native structure. The authors used Monte Carlo simulation of a C-alpha representation combined with an intramolecular energy potential that besides native interactions, also considers sequence-specific non-native interactions (i.e. non-native interactions that are encoded in the primary sequence) in the protein folding energetics. The outcome of this study provided further support for a functional role played by non-native interactions in the folding of knotted proteins by showing that (when compared with a purely native-centric potential) they enhance the probability to knot the polypeptide chain along the whole folding process (including conformations with a relatively low degree of structural consolidation), while assisting a knotting step based primarily on a direct threading movement of the C-terminus through loops formed by other parts of the chain. Similarly, a native centric potential complemented with (sequence-dependent) non-native interactions also leads to an enhancement of knotting frequency in association with a direct threading movement of the C-terminus for protein MJ0366^29^ (PDB ID: 2efv). The latter is the smallest knotted protein identified so far and features a shallow knot. Interestingly, a direct movement of the C-terminus was also observed in full atomistic Molecular Dynamics simulations of MJ0366 based on a classic force field^29^, reinforcing the idea that this is likely the mechanism that nature uses to tangle this protein. Overall, these results do suggest that non-native interactions play a functional role in the folding of knotted proteins by assisting (or even driving) a knotting step that is based on a direct movement of the protein’s C-terminus. Furthermore, they indicate that knotted conformations can form rather early in the folding pathway, in conformations which show a small degree of structural consolidation. Based on the analysis of folding trajectories obtained for MJ0366, Facciolli and co-workers suggested that non-native interactions promote the correct order of native contact formation, which is crucial for a protein to efficiently self-tie^29^. Indeed, it is widely accepted that the folding of knotted proteins stands out for being remarkably ordered^30^ (e.g. in the case of MJ0366 the two beta strands that exist in its native structure need to form the beta-sheet prior to any threading events^29^). At a microscopic level, an ordered folding process means that the establishment of native contacts must follow a specific sequence of events^8,20^. This ordered folding is the reason why knotted proteins are especially prone to backtracking, i.e., the formation, breaking and re-establishment of contacts that form untimely^31^. Molecular simulation studies based on native centric potentials appear to be less efficient in tying a knotted protein, presumably because they favor the premature (and likely untimely) establishment of interactions that are in close proximity in the native conformation. Possibly because of backtracking, the folding efficiency of these models is lower, and knotting occurs later in nearly native conformations, mainly through the mechanism of slipknotting^32,33^.

Since native interactions are critical for folding, and non-native interactions appear to be important for efficient knotting, it is important to understand their relative role, and this has been pointed out as a major question in a recent review of this research theme^34^. The results outlined above lead us to question if non-native interactions are strictly necessary to efficiently tangle a protein, or if it is possible that the correct order of contact formation can be assured by a specific set of native, but otherwise heterogeneous, interactions. This is the major question we propose to address in the present study, and in order to answer it we will use Monte Carlo simulations of lattice models. The reason is two-fold. Firstly, we are interested in determining the order of contact formation, and lattice models define unequivocally native and non-native contacts. Secondly, we are interested in exploring the folding and knotting behavior of proteinlike sequences with an intramolecular potential that treats identically the contribution of sequence-specific native and non-native interactions to the protein energy. Such an intramolecular potential has a long history in the study of protein folding^35–37^. Our results show that specific (non-local) non-native interactions indeed play a role in assisting the threading movement of the terminus in a particular ensemble of conformations with roughly half of the native contacts, but, more importantly, they also clearly show that there is a specific set of native interactions which are able to ensure optimal knotting performance by driving a folding pathway with minimal backtracking, which is populated by a knotted intermediate state that forms relatively early during folding.

## Models and methods

We use a simple lattice model in which amino acids are represented by beads of uniform size that occupy the vertices of a regular cubic lattice. To enforce the excluded volume constraint, only one bead is allowed per lattice site. Consecutive beads along the chain are connected by sticks (with length of one lattice spacing) that represent the peptide bond. Intramolecular contacts between two non-bonded beads occur if they are separated by one lattice spacing. Intramolecular contacts are classified either as native, if they are present in the native conformation, or non-native if they are established between any two beads that are not in contact in the native conformation.

To model protein energetics we consider the native centric Gō potential^27^, and two sequence-specific potentials: the one developed by Shakhnovich^38^, which we used in prior studies^36,37,39,40^, and a new native-centric potential with heterogeneous interactions. In all three cases, energy is measured in reduced units.

### The Gō potential

The total energy of a conformation defined by bead coordinates 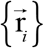 is given by

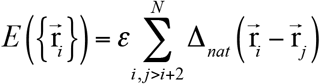

In the equation above, ε is a uniform intramolecular energy parameter (taken as −1 in this study), *N* is the chain length measured in number of beads, and the so-called contact function, Δ_nat_, is unity, if and only if, beads *i* and *j* are establishing a native contact.

### Sequence specific potentials

In the case of a sequence specific potential the total energy of a conformation is given by the Shakhnovich potential

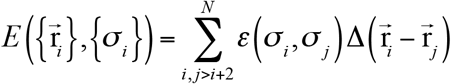

where {*σ_i_*} represents a specific sequence of amino acids that was designed to have minimum energy in the native conformation^41^, σ_i_ represents the chemical identity of bead *i*, and the intramolecular energy interaction parameters *ε*(*σ_i_, σ_j_*) are taken from the 20 × 20 Miyazawa-Jernigan interaction matrix^42^. Unlike for the Gō potential, the contact function, Δ, is unity if beads *i* and *j* are establishing an intramolecular contact, which means that both native and non-native interactions contribute to the energy of a conformation.

The difference between the Shakhnovich potential and the other sequence-specific potentials used in this study is that, in the latter, the contact function Δ, is unity if and only if beads *i* and *j* are establishing a native contact or some specific non-native contact(s).

The folding process is simulated by sampling canonically distributed conformational states generated using Metropolis Monte Carlo^43^ combined with replica exchange^44^. The conformational space was explored using a move set that comprised end-, corner-flip and crankshaft moves^39^. Following previous studies^9,13,20,33,45^ we used the fraction of native contacts, *Q*, as a reaction coordinate, i.e., as an indicator of folding progress, with *Q*~0 representing the denatured ensemble and *Q* = 1 representing the native state.

Simulations start from an unfolded conformation and terminate after 10^11^ Monte Carlo steps. The first half of the simulation is used to allow all replicas to relax to the equilibrium ensemble at their particular temperature, and relevant properties (*E, Q,* topological state, etc.) are then sampled during the second half. The topological state of a conformation (i.e. knotted or unknotted) was determined using the Koniaris–Muthukumar–Taylor (KMT) algorithm^25,46^. The Weighted Histogram Analysis Method (WHAM)^47^ was used to analyze the sampled data and produce maximum likelihood estimates of the density of states, from which expected values for thermodynamic properties were calculated as functions of temperature. WHAM was also used to project the density of states along the chosen reaction coordinate to obtain free energy and probability profiles along this coordinate. In particular, the specific heat capacity, *C_V_*, defined as *C_V_* = (< *E*^2^ > – < *E* >^2^)/ *T*^2^, was evaluated as a function of temperature, and the melting temperature, *T_m_*, was determined as the temperature at which *C_V_* reached its maximum value. The folding transition temperature, *T_f_*, was determined as the temperature at which the free energy profile had two minima of equal value, one near the unfolded state and the other near the native state. *T_m_* and *T_f_* should coincide for a two-state folding transition.

### Conditional probability contact maps

To obtain a microscopic description of the folding pathway we used conditional probability contact maps (CPCMs), i.e., *N* × *N* matrices where each entry represents the probability of a contact (native or non-native) being formed in an ensemble of conformations that share the same *Q* under thermal equilibrium at *T_m_*.

### Sequence design

To design protein sequences we performed design in sequence space^41^. Starting from a random sequence of *N* amino acids, an optimized (i.e. designed) proteinlike sequence was generated so that the energy of the native structure was minimal. This was achieved by means of Monte Carlo simulated annealing. Briefly, once an initial annealing temperature was selected, the chemical identity of two randomly selected beads was permutated at each MC step, this move being accepted or rejected according to the Metropolis criterion, and the temperature decreased according to a predefined annealing schedule. Further details on the design procedure can be found elsewhere^39^.

### Structural clustering

In order to determine the relevant conformational classes present in an ensemble of conformations we used the hierarchical clustering algorithm *jclust* available in the MMTSB tool set^48^. Since we are using a lattice model, clustering was based on contact map similarity. From each identified cluster we extracted the conformation that lies closest to the cluster’s centroid.

## Results

### Model systems and proteinlike sequences

In this work we consider a lattice protein with chain length *N* = 41 which was designed by hand to embed a trefoil knot (a 3_1_ knot) in its native structure^49^, and which, henceforth, will be named K3_1_ (Fig. 1A). The knotted core (KC) of K3_1_, i.e., the minimal segment of the chain that contains the knot, extends from beads 2 to 21. Since it suffices to remove two beads to untangle the protein, K3_1_ classifies as shallow.

**Figure 1.**
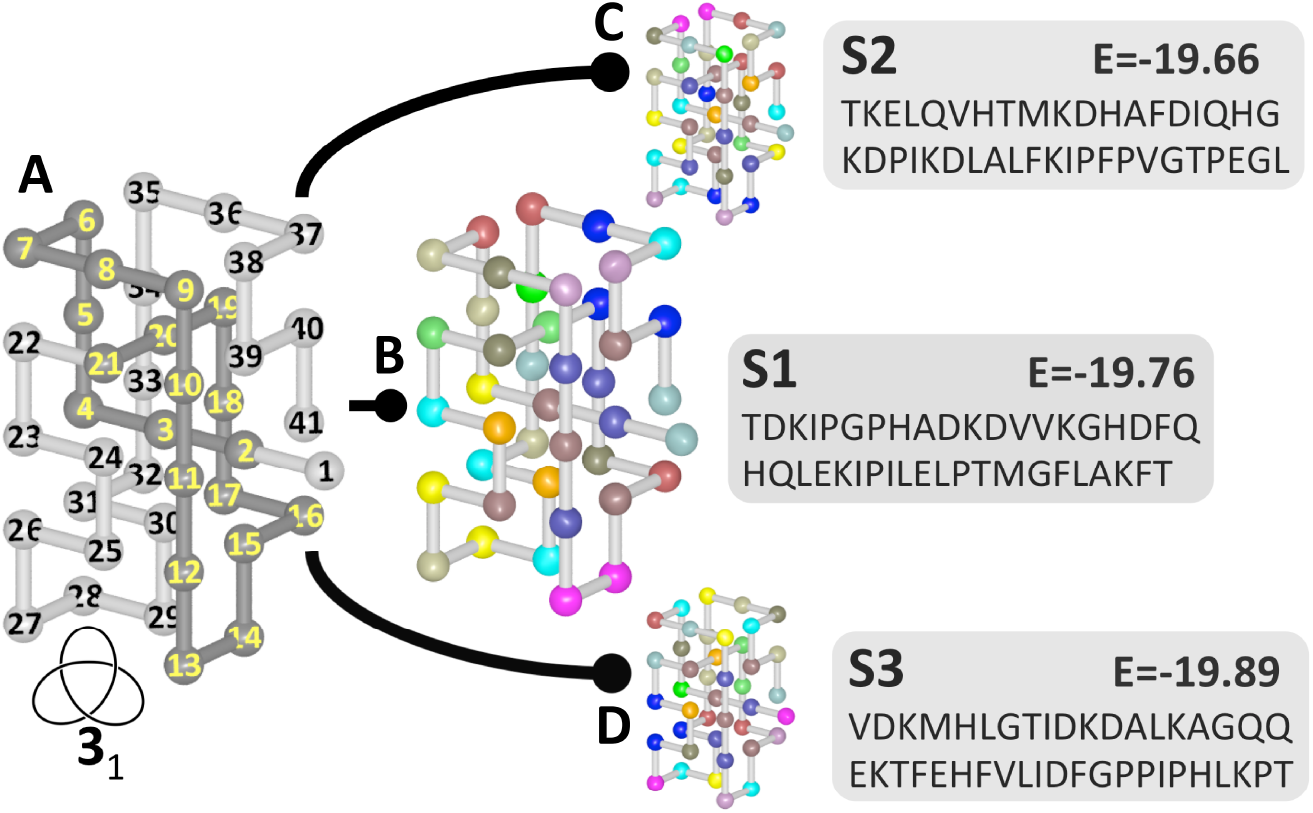
Model systems. (A) Native conformation of K3_1_ with knotted core (KC) highlighted in dark grey. Proteinlike sequences S1 (B), S2 (C), and S3 (D) and corresponding native energy.

Three proteinlike sequences, named S1, S2 and S3, were designed to have minimal (and similar) energy in the native structure of K3_1_ (Fig. 1B-D). It is important to mention that all sequences S1-S3 have the same number of different amino acids (i.e. all of them have 0 Cys (C), 1 Met (M), 3 Phe (F), 3 Ile (I), 4 Leu (L), 2 Val (V), 0 Trp (W), 0 Tyr (Y), 2 Ala (A), 3 Gly (G), 3 Thr (T), 0 Ser (S), 2 Gln (Q), 0 Asn (N), 2 Glu (E), 4 Asp (D), 3 His (H), 0 Arg (R), 5 Lys (K) and 4 Pro (P)), and only differ in the order by which the amino acids are assigned to the beads.

### Knotting transition

The major goal of this study is to determine if and how native and non-native interactions assist knotting. A possible strategy to address this problem is to perform a comparative analysis of the folding and knotting transitions of proteinlike sequences with different knotting performances, focusing on the behavior of native and and non-native interactions. We define the knotting probability, p_knot_, as being the fraction of knotted conformations in a given ensemble where all the conformations share the same Q. The dependence of p_knot_ on *Q* (evaluated at T_m_) indicates that for all the three sequences S1-S3 the knotting transition is a sigmoidal curve. However, S1 displays the earliest knotting transition (i.e. knotting occurs in less structurally consolidated conformations), S3 the latest (i.e. knotting occurs in more structurally consolidated conformations), and S2 exhibits an intermediate profile, becoming knotted in the vicinity of the transition driven by the Gō potential (Fig. 2A).

**Figure 2.**
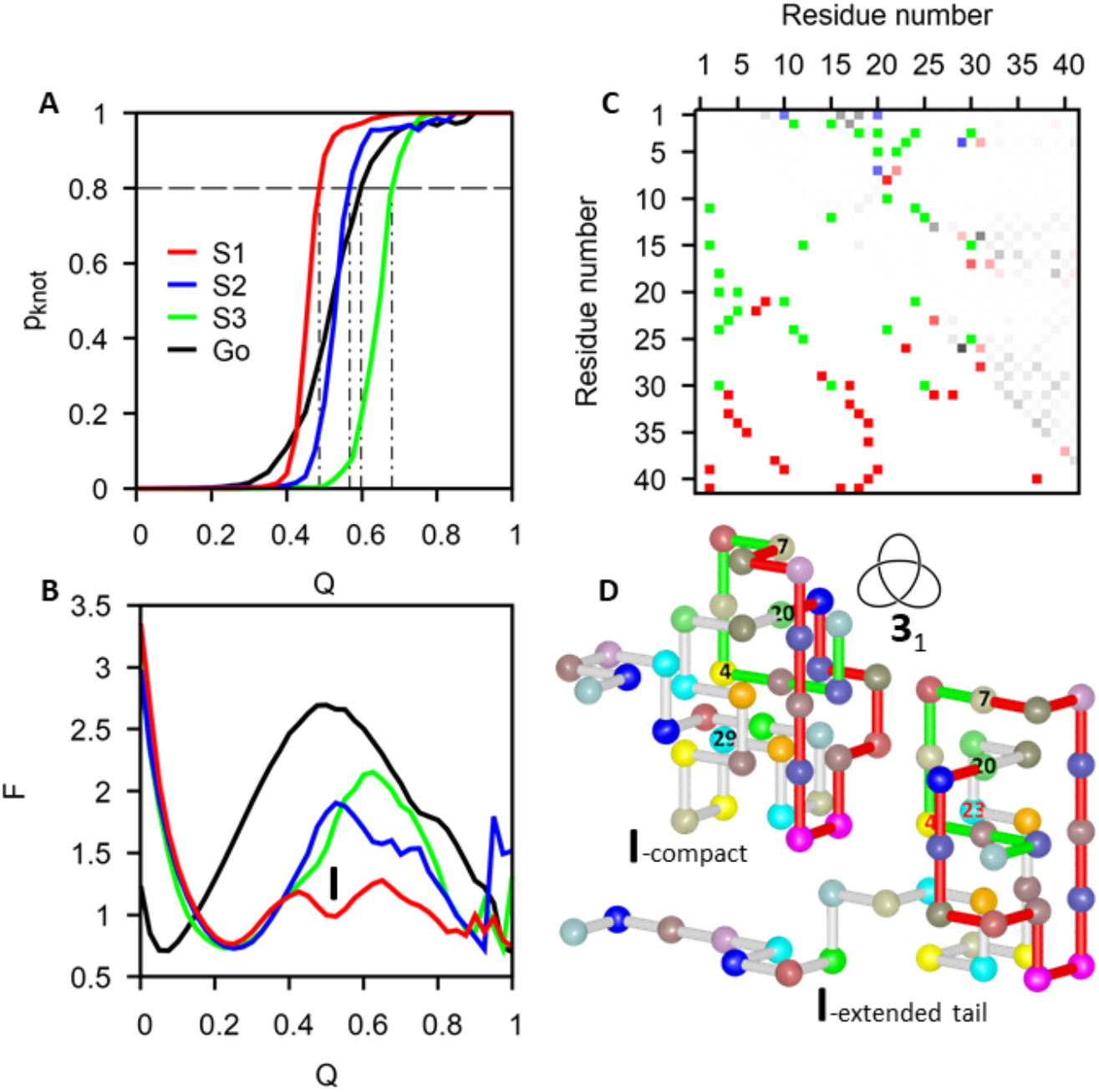
Knotting and folding transitions. (A) Knotting probability (p_knot_) as a function of the reaction coordinate (*Q*) for sequences S1-S3, and Gō, and (B) free energy profiles (to allow for a direct comparison, the native energy for the Gō potential was normalized to be the same as in S1). The intermediate state I populated by S1 at *Q* = 0.53. (C) The probability contact map for the intermediate state I (above the diagonal), and the native contact map, comprising 40 native contacts (below the diagonal). The 16 most probable native contacts in the intermediate are shown in shades of green, saturated proportionately to the contact probability, and the remaining native contacts are similarly shown in shades of red. The 4 most probably non-native contacts are shown in shades of blue, and the remaining non-native contacts in shades of grey. (D) Representative conformations (centroids) of the two (knotted) clusters found in the conformational ensemble of the intermediate state I. In both conformations, the knotting loop is highlighted in red, and the threading terminus is highlighted in green.

To get a more quantitative measure of how each protein sequence determines (by enhancing or suppressing) the probability to knot at a certain *Q*, we adopted the convention that the transition starts when p_knot_ = 0.2 (and term *Q_0.2_* the corresponding *Q* value), and ends when p_knot_=0.8 (and term *Q_0.8_* the corresponding *Q* value). We focus on *Q_0.8_*, which we take as indicator that the transition is nearly complete. We found that S1 requires 20 native contacts to reach a p_knot_ higher than 0.8 (*Q_0.8_* = 0.49), S2 requires 23 (*Q_0.8_* = 0.57), the Gō sequence 24 (*Q_0.8_* = 0.60), and S3 requires 28 native contacts (*Q_0.8_* = 0.68). The different knotting performance of sequences S1-S3 makes them ideal model systems to explore our strategy.

### Folding thermodynamics and intermediate states

Experimental results indicate that several knotted proteins with deep knots are prone to populate intermediate states^4,50–52^. Because knotting is entropically costly^53,54^, and intermediate states represent stabilized thermodynamic states, their formation along the folding pathway could serve the purpose of assisting the knotting step by decreasing the free energy barrier to knotting. In line with this hypothesis, and in order to determine what underlies the different knotting transitions of S1-S3, we investigated the existence intermediate states. The free energy profiles, i.e. the free energy projected along the reaction coordinate *Q*, show that the Gō sequence together with sequences S2 and S3 exhibit two-state transitions, but sequence S1, which is the one that shows the earliest knotting transition, populates an intermediate state (denoted by I in Fig. 2B) centered at *Q* = 0.53 (21 native contacts), i.e. slightly above the *Q_0.8_* = 0.49 found for this sequence. Since the native conformation is conserved in the four model systems, the formation of the intermediate state (I) must result from the specific amino acid interactions encoded in S1.

To structurally characterize the intermediate state we analyzed an ensemble of 12544 conformations with 21 native contacts (*Q* = 0.53) sampled by the replica having *T* nearest to *T_m_* (from above). In this conformational ensemble there are 11811 (94%) knotted conformations, which is consistent with S1 having *Q_0.8_* = 0.49. The probability contact map evaluated over the intermediate state ensemble (Fig. 2C) indicates that the intermediate state is stabilized by 20 contacts that form with highest probability, 16 of which are native (namely, 2-11, 2-15, 3-18, 3-20, 3-24, 3-30, 4-23, 5-20, 5-22, 10-21, 11-24, 12-15, 12-25, 15-30, 21-24 and 25-30), and 4 non-native (namely, 1-10, 1-20, 4-29 and 7-20).

By performing structural clustering over the ensemble of knotted conformations one is able to identify two clusters: a dominant cluster with 7118 (60%) elements, and a smaller cluster comprising 4693 (40%) elements. The dominant cluster is represented by a compact conformation, and the smaller cluster is represented by an extended C-tail conformation (Fig. 2D). Visual inspection shows that non-native contact (7-20) temporarily stabilizes the knotting loop in both conformations, while threading (the passage of the terminus through the knotting loop) is assisted by non-native contact (4-29) in the compact conformation, and by native contact (4-23) in the extended tail conformation.

### Contact formation, ordered folding and backtracking

The results reported in the previous section suggest that the mechanism behind the particular knotting transition of S1 is the formation of a knotted intermediate state that is energetically stabilized by 16 native interactions, and 4 non-native interactions. The question, then, is why does S1 populate this intermediate state while the other two sequences – which fold to the same native structure, have nearly the same native energy and same amount of each different amino acid in their chemical composition – do not? To address this problem we decided to explore the individual behavior of each native and nonnative contact by evaluating their probability profiles, i.e., their conditional probability of formation along the reaction coordinate *Q*. The rationale for doing so is the following. It is known that the folding of knotted proteins stands out for being remarkably ordered^29,30,55^.

At a microscopic level this means that the establishment of native contacts must follow a specific sequence of events. The need for an ordered contact formation is the reason why knotted proteins are especially prone to backtracking^20,32,56^, i.e., the formation, breaking and re-establishment of contacts that form untimely^57^. Our working hypothesis is that the level of backtracking is smaller in S1 than in the other 2 sequences, possibly because the intramolecular energies that were assigned to each native contact of S1 drive their formation in an order that is compatible with the constructive accumulation of structure giving rise to the observed intermediate state.

We find that contact probability profiles come out in different shapes but, for native contacts, it is possible to distinguish two major profile types: a sigmoidal shaped probability profile (Fig. 3A), and an irregularly shaped profile, exhibiting humps, bumps and plateaus, which are indication of backtracking (Fig. 3B). Non-native contacts have somewhat bell shaped probability profiles, which often also exhibit considerable irregularities (Fig. 3C). Interestingly, the vast majority (35 out of 40) of native contacts in sequence S3, display this type of irregular profile (Fig. 3B), while for S1 it is possible to find 10 native contacts with a clear sigmoidal profile out of the 16 contacts that precede the knotting transition (Fig. 3A). These results thus indicate that sequence S1 is the one that exhibits a smaller level of backtracking, which allows the formation of a knotted intermediate that is on-pathway to the native state. The formation of this intermediate state is concomitant with an expressive lowering of the major free energy barrier (Fig. 2B), and thermodynamic stabilization of the full folding transition.

**Figure 3.**
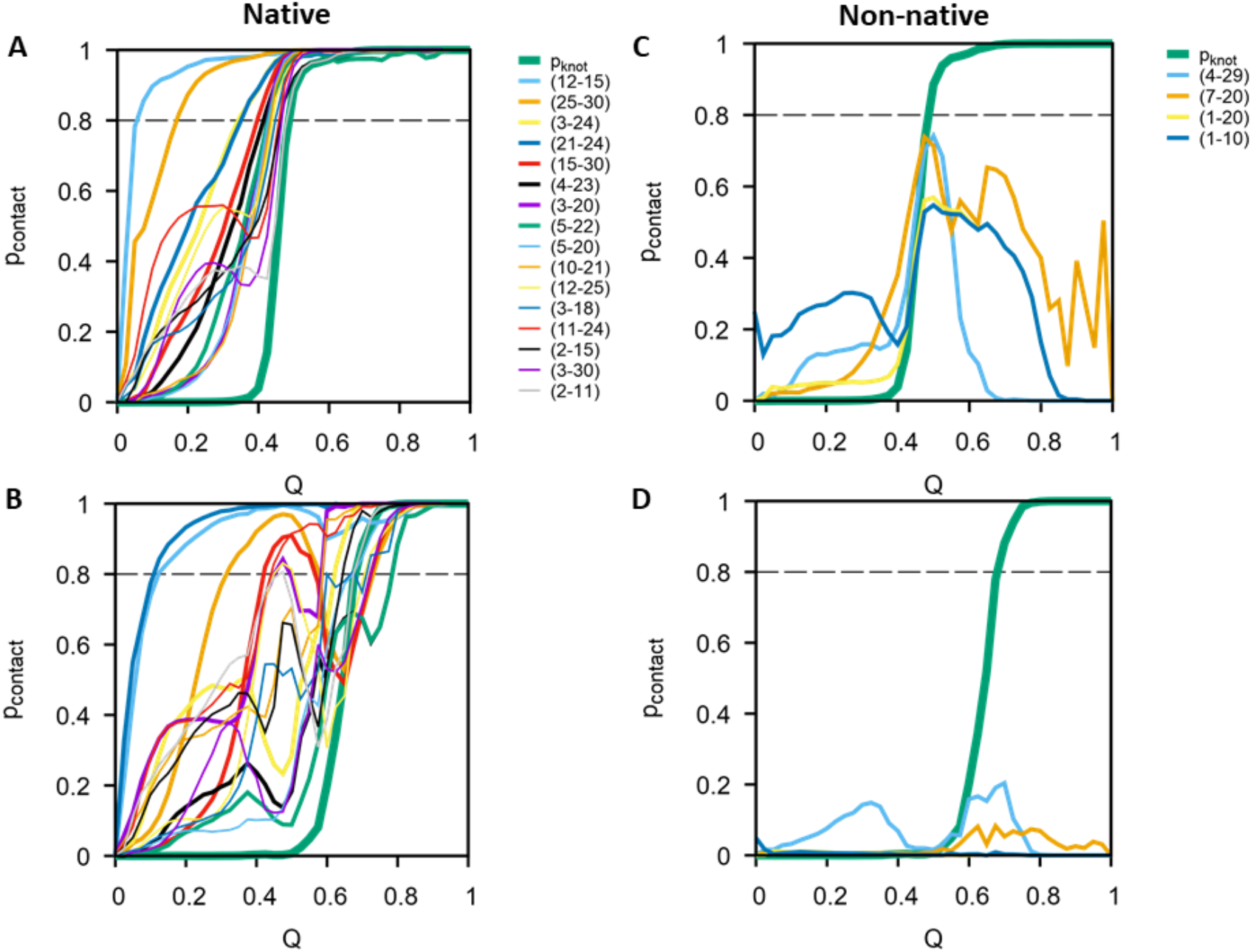
Contact probability profiles. Probability profiles of native contacts start at p = 0 (*Q* = 0) and end at p = 1 (*Q* = 1), whereas the probability profiles of non-native contacts may start at any value for *Q* = 0 and must end at p = 0 for *Q* = 1. (A) Profiles of the 16 native contacts that precede knot formation in sequence S1.These are the same contacts found to be most probable in this sequence’s intermediate state (SI Table I). (B) Profiles of the same 16 native contacts in the folding pathway of S3. (C) Profiles of the 4 most probable non-local non-native contacts identified in the intermediate state of S1 (SI Table I). (D) Profiles of the same 4 non-native contacts in the folding pathway of S3.

Similarly to the knotting transition, the value of the reaction coordinate at which a native contact’s probability profile crosses the 0.8 probability threshold is named the native contact’s *Q_0.8_* (i.e. for all values of *Q ≥ Q_0.8_*, p_contact_ ≥ 0.8), and is an indicator of the stage at which the native contact becomes locked (or permanently formed) during the folding process. The sequence in which the native contacts get locked is a possible way to characterize the folding pathway, and this sequence can be traced by sorting the native contacts by increasing order of their *Q_0.8_*. We find that in the case of S1, 16 native contacts have *Q_0.8_* lower than the *Q_0.8_* = 0.49 of its knotting transition, and that these are exactly the same native contacts identified as most probable in the intermediate state I populated by this sequence.

Perhaps not surprisingly, when, for the native contacts of S1, we plot the (contact energy, *Q_0.8_*) pairs we find a statistically significant trend [for S1 (S3): linear term has t = 4.171 (5.469), null hypothesis p = 0.00017 (3.05×10^-6^)] between *Q_0.8_* and the contact’s intramolecular energy (SI: Fig. 1A, D), with the more stable native contacts tending to form before the less stable ones in less consolidated conformations. This means that, in principle, when designing a knotted protein, the sequence of native contact formation can be tuned by changing the relative contact energies of the native contacts.

For non-native contacts, the contact probability profile enables calculation of two indicators: p_max_ which is the maximum value reached by the conditional contact probability, and *Q_max_* which is the value of the reaction coordinate at which p_max_ occurs.

Unlike Q_0.8_ for native contacts, for non-native contacts, *Q_max_*, which indicates the stage in the folding pathway at which the contact is most relevant, is not significantly influenced by the contact’s intramolecular energy [for S1 (S3): linear term has t=−0.461 (−0.758), null hypothesis p = 0.645 (0.449)] (SI: Fig. 1B, E). The stage at which a non-native contact becomes relevant during the folding process can thus only be a consequence of geometric and excluded volume constraints that result from already formed native contacts. However, when these constraints allow the formation of a non-native contact, its probability is strongly influenced by the contact’s intramolecular energy [for S1 (S3): linear term has t = – 19.399 (−17.611), null hypothesis p < 2×10^-16^ (2×10^-16^), quadratic term has t = −2.222 (−1.351), null hypothesis p = 0.0269 (0.178)] (SI: Fig. 1C, F) and the level of assistance that a non-native contact may give to the stabilization of temporary intermediate conformations is, consequently, strongly dependent on its contact energy.

### Role of native and non-native interactions

The results reported in the previous sections indicate that the 16 native interactions that are the first to lock in the folding of S1 may, either alone, or together with specific non-native interactions, be responsible for the knotting performance of S1. However, it is possible that not all of the native interactions are strictly necessary for knotting. Since the ensemble representative of the intermediate state (I) populated by S1 contains knotted and unknotted conformations, we evaluated the frequency of native and non-native contacts, as well as the contact probability curves in each sub-ensemble. We find 5 non-local, non-native contacts (namely, 4-29, 1-10, 14-31, 1-20, 7-20) that are potentially important for knotting because they are clearly more frequent in the sub-ensemble of knotted conformations (SI: Table 1 and SI: Fig. 4A-E). On the other hand, we also identified 4 native contacts (namely, 25-30, 12-15, 4-23, 5-22, and 25-30) that may not be crucial for knotting, exhibiting similar probabilities in both sub-ensembles (SI Table I and SI Fig. 4F-I). In order to test these hypotheses we conducted several folding simulations of K3_1_ under different interaction potentials that consider only native interactions, or selected combinations of native and non-native interactions (Table 1).

**Table 1.** Intermolecular potentials used in this study. The 12 native interactions used in ss12 exclude native interactions 25-30, 12-15, 4-23 and 5-22 out of the 16 that are the first to lock in the folding of sequence S1.

| Potential | Native inter.<br>(from S1) | Native inter.<br>(homogeneous) | Non-native inter. |
| --- | --- | --- | --- |
| ss40 | 40 | 0 | 0 |
| ss40<sup>+</sup> | 40 | 0 | 4-29 or 7-20 |
| ss40<sup>+2</sup> | 40 | 0 | 4-29 + 7-20 |
| ss16 | 16 | 24 | 0 |
| ss16<sup>+</sup> | 16 | 24 | 4-29 or 7-20 |
| ss16<sup>+2</sup> | 16 | 24 | 4-29 + 7-20 |
| ss16<sup>0</sup> | 16 | 0 | 0 |
| ss12<sup>+5</sup> | 12 | 27 | 4-29 + 1-10 + 14-31 + 1-20 + 7-20 |
| ss12<sup>0</sup> | 12 | 0 | 0 |

We start by assessing the role of native interactions in knotting by considering two intramolecular potentials. The first, which we term sequence-specific 40 (ss40), takes into account the native interactions of sequence S1, and ascribes null energies to all non-native contacts. The second, which we term ss16, conserves the interaction parameters of S1 for the 16 native contacts found in the intermediate state, and combines them with homogeneous energies for the remaining 24 native contacts (so that the native state’s energy of sequence S1 is preserved), and ascribes null energy to all non-native contacts.

The knotting probability profiles of K3_1_ obtained under these two potentials are qualitatively similar, and show that the set of 16 native interactions are able to efficiently knot the chain. These two p_knot_ profiles, however, lag behind the p_knot_ profile of S1 between *Q* = 0.45 and *Q* = 0.53 and, hence, the knotting enhancement observed in S1 in this interval of the reaction coordinate cannot be explained by native contacts alone (Fig. 4A).

**Figure 4.**
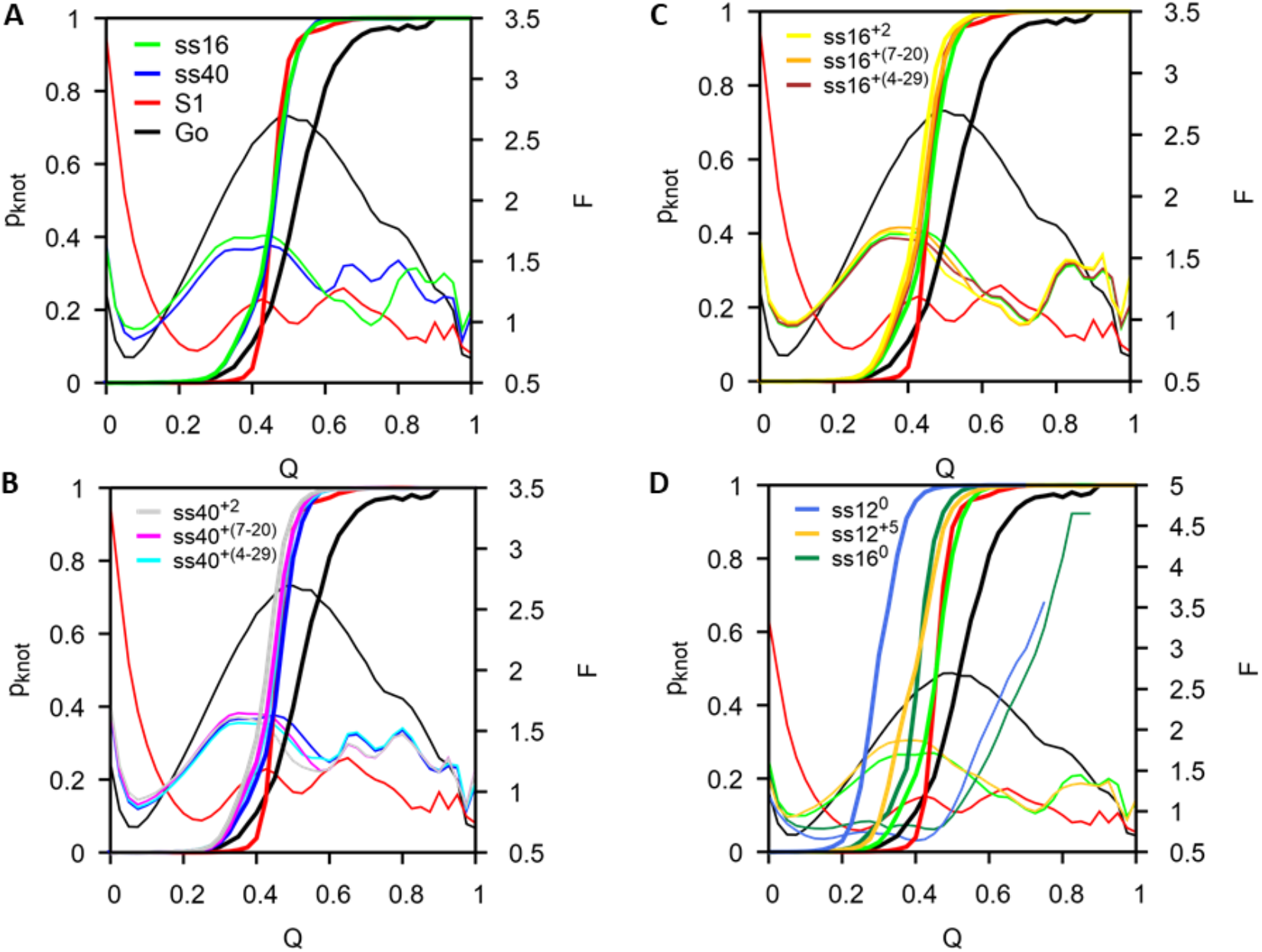
Role of native and non-native interactions in knotting. (A) Knotting probability and free energy profiles for potentials ss40 and ss16. (B) Profiles for potentials ss40^+(4-29)^, ss40^+(7-20)^ and ss40^+2^. (C) Profiles for potentials ss16^+(4-29)^, ss16^+(7-20)^ and ss16^+2^. (D) Profiles for potentials ss16^0^, ss12^+5^ and ss12^0^.

In fact, when, in addition to the native interactions, we consider non-native interactions (e.g. 4-29, or 7-20), we obtain the p_knot_ profiles labeled ss40^+(4-29)^ and ss40^+(7-20)^ (Fig. 4B), or ss16^+(4-29)^ and ss16^+(7-20)^ (Fig. 4C), which nearly coincide with the S1 profile in this interval. Adding both non-native interactions to ss40 (ss40^+2^) (Fig. 4B), or ss16 (ss16^+2^) (Fig. 4C), leads to even earlier knotting profiles.

The free energy profiles of ss40, ss16 show two major differences from that of sequence S1. In both cases, one observes the formation of a novel, more structurally consolidated (*Q* ~ 0.7), intermediate state (Fig. 4A). Structural clustering analysis over an ensemble of 16423 conformations with *Q* = 0.7 populated by ss16 reveals the existence of 5 clusters of knotted conformations (instead of only 2 detected for the intermediate state populated by S1) (SI Fig. 2). Secondly, and more importantly, the transition state (TS) that precedes the formation of the intermediate state (located at *Q* = 0.43 for S1 and *Q* = 0.45 for ss40) is substantially more stabilized in S1 than in ss40, but only 15% of its conformations are knotted. Under both potentials the thermodynamic stabilization of the TS is dominated by the energy component of the free energy (U_TS_=-14.95 for S1, and U_TS_=-11.41 for ss40), but in the case of S1 the latter represents 79% of the free energy, and only 59% in the case of ss40. The energy associated with non-native interactions represents 30% of the total energy of the TS preceding the formation of the intermediate of S1, which allows us to conclude that non-native interactions are responsible for its higher thermodynamic stability. Interestingly, a similar analysis carried out for the intermediate state of S1 reveals that its thermodynamic stabilization is also dominated by the energetic contribution, but the energy of non-native interactions represents only 12% of the total energy.

To test if the 16 native interactions were sufficient to drive the knotting transition, we considered a potential (ss16^0^) in which the 16 native contacts kept their S1 energies, and all other contacts (native and non-native) were given null energies. We find that the 16 native interactions are indeed sufficient to knot the chain through the formation of conformations with an even smaller degree of structural consolidation (Fig. 4D). It is interesting to note that the knotting mechanism changes, however. The contact that forms concomitantly with knotting is now (10-21) instead of (2-11) (SI Fig. 3A), which indicates that knotting is now completed through a mouse-trap motion^10,29,59^ rather than direct threading of the terminus. The considered interactions, however, are not sufficient to fold the protein, which requires the contribution of all native interactions (Fig. 4D).

Are all of these 16 native interactions required for knotting? In order to answer this question, we considered yet another intermolecular potential that takes into account the S1 energies of the 12 native contacts and 5 non-native contacts (that are more relevant in the sub-ensemble of knotted conformations), ascribes null energies to all other non-native contacts, and uniform energies to the remain 27 native contacts so that the energy of the native structure is preserved. Remarkably, this potential (termed ss12^+5^) leads to an even earlier knotting transition, showing that the corresponding interplay of native and nonnative interactions is optimal to fold this particular lattice protein. It is worth mentioning that the addition of non-native interactions (in this case and the other cases analyzed before) does not change the free energy profiles, indicating that non-native interactions do not play a thermodynamic role in assisting the knotting step (Fig. 4D). Instead, the considered specific non-native interactions play a structural or kinetic role instead, by temporarily stabilizing the knotting loop or assisting the threading step (SI Fig. 3B).

A modification of this potential that switches-off the 27 native interactions and non-native interactions (ss12^0^) leads to the earliest knotting transition observed in this study, but the protein is not able to fold (Fig. 4D), in line with what is observed for ss16^0^.

## Conclusions

The role of non-native interactions in the folding of knotted proteins has been pointed out as an important fundamental question in the field of protein folding. Results reported in the literature are consistent with the idea that, independently of the sampling scheme used to probe the conformational space (Monte Carlo or Molecular Dynamics), and of the resolution of protein representation adopted in simulations (from single bead to full atomistic), non-native interactions are critical to knot the chain, at least when the latter involves a single threading event. While the mechanistic details according to which they operate remain elusive, it has been suggested that non-native interactions assist knotting by driving the correct order of contact formation thus avoiding backtracking.

Here we explored this hypothesis in the context of a lattice model. In lattice models, native and non-native contacts are defined unequivocally and, therefore, lattice models offer an appropriate framework to investigate the order of contact formation.

By using a sequence specific potential that naturally incorporates the participation of nonnative interactions in protein energetics, we were able to find nativelike sequences with different knotting performances. In particular, when gauged against a native-centric Gō model, we found that some sequences are more efficient and knot in conformations with a small degree of structural consolidation, while others need a higher degree of structural consolidation to knot.

By making an in depth comparative study, which comprised a thermodynamics analysis, and a microscopic investigation at the level of each individual contact, we were able to find efficient knotting does not require non-native interactions. Notwithstanding, we were able to identify specific non-native interactions that play a functional role in the knotting process by acting like a scaffold that temporarily stabilizes the knotting loop, or assists the threading step. Efficient knotting is, on the other hand, driven by a critical set of specific native interactions, which do not backtrack, and contribute to stabilize an intermediate state that decreases the entropic cost to knotting. The reason behind the ‘ideal’ behavior of these native interactions relies, at least to a significant extent, on their relative energies (with the more stable native interactions tending to consolidate first), and how these energies are distributed within the native fold to minimize energetic and topological bottlenecks, which are being particularly important in the case of knotted proteins. The fact that efficient knotting only requires 12 out of 40 native interactions (which represents a reduction of 30% of the original amino acid alphabet) is interesting *per se,* and suggests that while energetic heterogeneity is important to assure a well defined, and correct order of contact formation, a 20 amino acid alphabet may not be necessary to efficiently tie a polypeptide chain. In light of these results the principle of minimal frustration might thus be restated for the particular case of knotted proteins as: knotted proteins evolve towards primary structures that underlie sigmoidal shaped contact probability profiles for their native contacts. A relevant question is then, what is the minimal size of the amino acid alphabet that efficiently ties a protein chain?

In this study we conducted replica-exchange Monte Carlo simulations, because our major goal was to study equilibrium thermodynamics. Another interesting question is to investigate the performance of the utilized potentials in simulations at fixed temperature to investigate their kinetic performance. In particular, which potential knots the chain faster?

It is also pertinent to ask whether or not similar results might be obtained in the scope of an off-lattice model or if, on the other hand, non-native interactions are indeed critical to drive an efficient knotting mechanism in off-lattice models, and perhaps in real world natural proteins. One should bear in mind that conformational entropy is measured more accurately off-lattice; likewise, the energetic stabilization associated with non-native interactions at the level of the unfolded or partially folded states, may become only visible off-lattice.

Furthermore, it is also possible that non-native interactions may play a structural role (e.g. by enlarging the knotting loop to avoid steric clashes), which is smeared out by the discrete nature of the lattice representation.

The questions outlined above will be the focus of future work.

## Author contributions

PFNF designed the research. JNCE performed the calculations. JNCE and PFNF analyzed the data. PFNF and JNCE wrote the paper.

## Acknowledgments

Work supported by UID/MULTI/04046/2020 Research Unit grant from FCT, Portugal (through BioISI). JNCE acknowledges financial support from FCT, Portugal, through PhD grant SFRH/BD/144345/2019. PFNF would like to thank FCT, for financial support through grant number PTDC/FIS-OUT/28210/2017. PFNF and JNCE would like to acknowledge the contribution of the COST Action CA17139.

